# A high-efficiency scar-free genome editing toolkit for *Acinetobacter baumannii*

**DOI:** 10.1101/2022.04.15.488490

**Authors:** Rubén de Dios, Kavita Gadar, Ronan R McCarthy

## Abstract

**Background:** The current mutagenesis tools for *Acinetobacter baumannii* leave selection markers or residual sequences behind, or involve tedious counterselection and screening steps. Furthermore, they are usually adapted for model strains, rather than to multidrug resistant (MDR) clinical isolates.

**Objectives:** To develop a scar-free genome editing tool suitable for chromosomal and plasmid modifications in MDR *A. baumannii* AB5075.

**Methods:** We prove the efficiency of our adapted genome editing system by deleting the multidrug efflux pumps *craA* and *cmlA5*, as well as curing plasmid p1AB5075. We then characterised the antibiotic sensitivity phenotype of the mutants compared to the wild type for chloramphenicol, tobramycin and amikacin by disc diffusion assays and determined their minimum inhibitory concentration for each strain.

**Results:** We successfully adapted the genome editing protocol to *A. baumannii* AB5075, achieving a double recombination frequency close to 100% and securing the construction of a mutant within 10 work days. Furthermore, we show that the Δ*craA* has a strong sensitivity to chloramphenicol, tobramycin and amikacin, whereas the Δ*cmlA5* mutant does not show a significant decrease in viability for the antibiotics tested. On the other hand, the removal of p1AB5075 produced an increased sensitivity to tobramycin and amikacin.

**Conclusion:** We have adapted a highly efficient genome editing tool for *A. baumannii* and proved that *craA* has a broader substrate range than previously thought. On the other hand, whereas *cmlA5* is annotated as a chloramphenicol efflux pump and is encoded within an aminoglycoside resistance island, it does not provide resistance to any of those compounds.

## Introduction

*A. baumannii* is an aerobic Gram-negative bacterium that is widespread in the environment and inhabits different niches. However, it can also be an opportunistic pathogen that infects immunocompromised patients.^1,2^ Nowadays, it is estimated that up to 10% of nosocomial infections in the United States and 2% in Europe are caused by this pathogen, with these frequencies almost doubling in Asia and the Middle East. Furthermore, around 45% of *A. baumannii* isolates in global terms exhibit multi-drug resistance (MDR, i.e. resistance to at least 3 classes of antibiotics), with local rates rocketing to 70% in Latin America and the Middle East.^1,3-5^ Due to this, *A. baumannii* has been included among the most concerning MDR pathogens under the acronym *ESKAPE* (*Enterococcus faecium, Staphylococcus aureus, Klebsiella pneumoniae, A. baumannii, Pseudomonas aeruginosa* and *Enterobacter spp*.).^6^ Moreover, a World Health Organisation (WHO) report highlighted carbapenem-resistant *A. baumannii* as a priority pathogen, for which novel therapeutic approaches urgently need to be developed.^7^

The recalcitrance of this species to treatment is due to its capacity for resistance and persistence,^1^ aided by its multiple MDR mechanisms. These include the cell envelope as a barrier, multi-drug efflux systems and mutations in genes coding for porins and antibiotic targets (e.g. ribosomal proteins, penicillin binding proteins, DNA replication enzymes and the lipid A biosynthetic pathway), as well as enzymes that degrade/inactivate antibiotics.^2^ Oftentimes, these features can spread among the population through mobile genetic elements and the ability of *A. baumannii* to be naturally competent.^2,8-10^

With technological advances, genome editing tools have evolved, allowing precise genome editing (i.e. insertions and deletions), from a single nucleotide to dozens of kilobases. However, this progress is often uneven, with tools being biasedly developed for a few well established model organisms. In the case of *A. baumannii*, many simple targeted genetic tools have been adapted for their use in model strains of this pathogen. These tools go from gene disruption by plasmid insertion in a single recombination event to mutation by antibiotic resistance marker insertion.^11^ Next-step strategies include recombineering-based gene disruption followed by removal of the selection marker by site-specific recombination, allowing the use of the same marker for subsequent rounds of mutation to construct multiple mutants.^12^ Even more refined, some protocols allow scar-less gene modification by double recombination aided by a counterselectable marker.^13^ Moreover, after the bloom of clustered regularly interspaced short palindromic repeats (CRISPR)-Cas systems as a molecular biology tool, a CRISPR interference (CRISPRi) kit has been developed for *A. baumannii* that allows knocking down the expression of both essential and non-essential genes.^14^ However, depending on the purpose they are intended for, these genetic editing methods can have some limitations. Gene disruption is not always desirable due to the limited amount of selection markers available and possible polar effects within operons. Strategies including marker removal are usually based on site-specific recombinases that leave a scar in the genome.^12,15^ However this recombinogenic sequence may cause genomic instability after successive rounds of mutation.^16^ These drawbacks can be prevented by counterselection-mediated scar-free strategies, which allow more complex genome manipulation (i.e. targeted point mutations, domain truncations, allele exchange, deletion of whole clusters), but counterselection often requires passaging under pressing selection and tedious screening for clones that underwent a second recombination event. Furthermore, the current tools are mainly developed for model *A. baumannii* strains, which can be less representative as compared to the prevalent clinical isolates. Besides, an extra limitation appears when applying these tools to MDR *A. baumannii* strains due to the little availability of selection markers.

In our efforts to implement state-of-the-art methodologies for standardisation of genome editing in non-model MDR *A. baumannii* strains, we have adapted an accelerated highly efficient SceI-based mutagenesis method,^17-20^ developed and optimised for *Pseudomonas putida*,^16,21^ to MDR *A. baumannii* AB5075.^3^ For this, we have modified the two plasmids used in this system with selectable markers that can be used in this strain and subsequently adapted the protocol pipeline. As a proof of concept, we have constructed an in-frame deletion mutant in *craA*, a gene encoding a dedicated chloramphenicol-specific efflux pump. Afterwards, we have attempted to address the function of *cmlA5*, a putative plasmid-borne chloramphenicol efflux pump coding gene inferred from homology, by comparison with the *craA* mutant. As a result, we have validated the utility of this system for scar-free chromosomal and plasmid editing in *A. baumannii* AB5075.

## Materials and Methods

### Bacterial strains and culture media

*A. baumannii* AB5075 (VIR-O colony morphotype),^3,22,23^ its derivate mutants and *E. coli* host strains (DH5□ and DH5□λ*pir*) were routinely grown in liquid or solid LB (Miller) at 37 ºC (180 rpm or static, respectively).^16,24^ When necessary, LB was supplemented with kanamycin (25 mg/L), ampicillin (100 mg/L), apramycin (60 mg/L for *E. coli*, 200 mg/L for *A. baumannii*), tetracycline (5 mg/L) or tellurite (6 mg/L for *E. coli*, 30 mg/L for *A. baumannii*). A summary of strains used in this work is shown in Supplementary Table S1.

### Plasmid construction

A list of plasmids and primer sequences used in this work can be found in Supplementary Table S1. All plasmid derivatives were constructed using standard restriction-based molecular cloning.

pEMG-Tel (pEMGT) was constructed by cloning a DNA fragment from pMo130-TelR (Addgene, #50799) (bearing the Tel resistance marker) digested with SmaI in pEMG cut with AflIII and blunted with Klenow.^13,16^ For construction of pSW-Apr and pSW-Tc, PCR fragments amplified from pFLAG-attP (Addgene, #110095) with primers Apr fw/Apr rv and from pSEVA524 with primers tetA fw/tetA rv,^25^ respectively, using Q5 High-Fidelity Master Mix (New England Biolabs) were cloned into pSW-I digested with ScaI.^16^

For in-frame deletion of *craA* (ABUW_0337) and *cmlA5* (ABUW_4059) pEMGT-*craA* and pEMGT-*cmlA5* were constructed. For pEMGT-craA, 1 kb upstream and downstream homologous regions were amplified from purified AB5075 genomic DNA with primers craA up fw/craA up rv and craA down fw/craA down rv, respectively, and assembled together by joining PCR. The same procedure was followed for assembly of the *cmlA5* deletion construct using primer pairs cmlA5 up fw/cmlA5 up rv and cmlA5 down fw/cmlA5 down rv. Both constructs were cloned into pEMGT digested with SmaI.

All plasmid derivatives were checked by colony PCR using DreamTaq Green PCR Master Mix (ThermoFisher), restriction patterns and eventually by Sanger sequencing.

### Triparental mating

For transfer of plasmid DNA into *A. baumannii* AB5075 and derivative strains, a standard triparental mating protocol was followed, using pRK2013 (in a DH5□ host) as helper plasmid and a DH5□ or a DH5□λ*pir* donor bearing the plasmid of interest.^26^ For each mating, 500 μl of overnight cultures of the respective receptor, helper and donor strains were mixed. Cells were pelleted by centrifugation and washed 2 times with fresh LB medium. The final cell pellet was resuspended in 40 μl of LB, spotted on a plain LB agar plate and left to air-dry. After that, the biomass patch was incubated at 37 ºC for 4 h. The biomass was then resuspended in 1 ml of LB and serial dilutions were spread on the respective selective media and on plain LB plates to assess viability and incubated at 37 ºC overnight. Selection and marker exchange were checked by multiple streaking on different selective LB plates supplemented with ampicillin (to select AB5075 against *E. coli* strains) and the suitable selective agent according to the resistance marker transferred plasmid. When necessary, DNA deletions in the receptor strain were assessed by colony PCR and eventual Sanger sequencing from PCR-amplified genomic DNA. Conjugation frequency was calculated as the number of transconjugant colonies divided by the number of viable cells.

### Antibiotic disc diffusion assay

Antibiotic sensitivity assays were performed in cation-adjusted (CaCl_2_ 10 mM, MgCl_2_ 5 mM) Mueller-Hinton (pH 7.4) medium (CAMH, Sigma-Aldrich). Overnight cultures of *A. baumannii* AB5075 or the respective mutant derivatives were diluted to 0.5 McFarland units in CAMH broth and spread with a cotton swab on CAMH agar plates. When plates were dry, chloramphenicol, amikacin or tobramycin discs (Oxoid) were placed in the middle of the CAMH agar plate. Plates were incubated at 37 ºC for 24 h before measuring the diameter of the inhibition zone. Results are shown as averages of 3 biological replicates.

### Minimum inhibitory concentration determination

Saturated overnight cultures were diluted in PBS (phosphate-buffered saline) to get an OD_600_ 0.2. The bacterial solutions were centrifuged at 6,000 rpm for 5 minutes and they were then washed 3 times in PBS. Biomass was then resuspended in 1.2 ml of CAMH broth. The dilution range of the antibiotic was prepared from a 50 mg/ml stock solution in CAMH broth. The starting concentration of antibiotic in the range of dilution was 2500 μg/ml and was then diluted 2-fold over 9 additional serial dilutions in CAMH broth.

In order of highest to lowest dilution, 100 μl of the antibiotic solution was added to each well on a 96-well plate. Next, 100 μl of the cell suspension were added to each well. As control, 100 μl of sterile CAMH broth plus100 μl of the bacterial solution and 200 μl of sterile CAMHB were tested. The 96-well plate was then incubated at 37 °C, 200 rpm. Final OD_600_ was measured after 16 h using a Clariostar Plus microplate reader (BMG LabTech). MICs were assessed by visual examination, defining it as the lowest antibiotic concentration that led to absence of visible bacteria growth.

## Results and discussion

### Rationale of the strategy

To adapt an efficient genome editing system for MDR *A. baumannii* AB5075, we built our strategy on that by E. Martínez-García and V. de Lorenzo for *P. putida*,^16^ further optimised to an accelerated version at the Nickel laboratory.^16,21^ To perform this strategy, plasmid pEMG and pSW-I need to be used.^16^ pEMG is a cloning suicide vector bearing two target sites for the endonuclease SceI flanking its polylinker. Once the homologous regions flanking the desired modification are cloned into pEMG, the resulting plasmid is transferred to the target strain and the integration in the genome is selected. Subsequently, the broad-host range pSW-I plasmid, the SceI coding gene under an inducible XylS-dependent promoter, is introduced in the co-integrate strain. Inducing the expression of *sceI* would trigger the double-strand break in the genome that would eventually be repaired by homologous recombination, generating the reversion to the parental strain genotype or the desired mutation. Apart from improvements to make the screening more efficient, Wirth *et al*. introduced on-plate induction of *sceI* expression,^21^ reducing the second recombination to one plasmid transfer and selection step.

In the case of *A. baumannii* AB5075, one of the disadvantages for its genetic manipulation is its resistance to most selectable markers, including those in pEMG and pSW-I. Hence, we tackled the construction of a pEMG derivative bearing a tellurite resistance cassette as well as its orginal kanamycin resistance gene. As a result, we obtained plasmid pEMG-TelR, abbreviated pEMGT (Figure 1). For the second part of the strategy, we produced two variants of the pSW-I plasmid, each bearing either an apramycin resistance marker or a tetracycline resistance gene, namely pSW-Apr (Figure 1) and pSW-Tc, respectively. These plasmids would serve as a platform for *A. baumannii* genome editing. To validate the method and demonstrate its versatility, we attempted the construction of scar-free mutants in the chromosome-encoded gene *craA* and the plasmid-borne gene *cmlA5*. Whereas *craA* (identified in AB5075 by sequence similarity to the *craA* orthologue characterised in *A. baumannii* ATCC 17978) is an efflux pump previously thought to be specific to chloramphenicol,^27,28^ but recently shown to have a broader substrate range,^29^ *cmlA5* is a putative chloramphenicol efflux pump inferred from homology and encoded within an aminoglycoside resistance island.^22^

**Figure 1.**
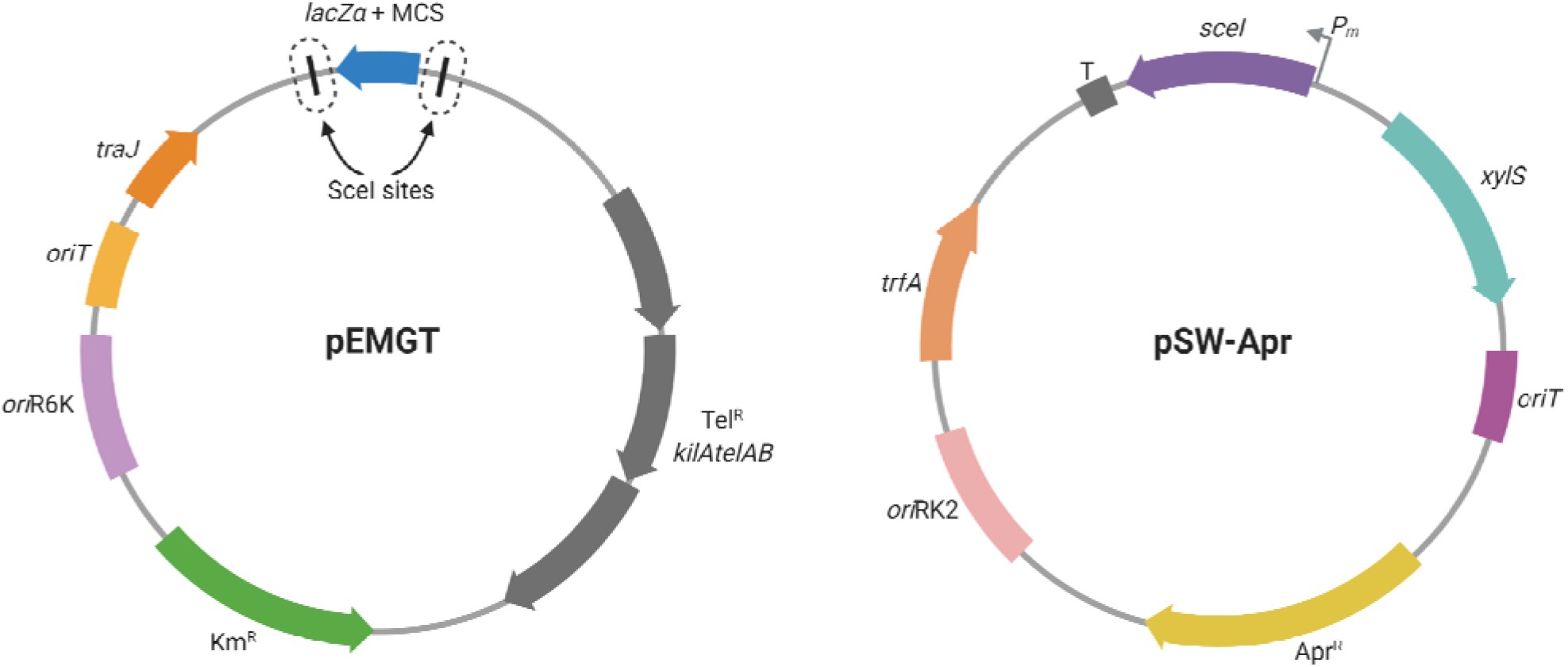
Schematic representation of plasmids pEMGT and pSW-Apr. All relevant features borne in each plasmid are presented and named. SceI target sites in pEMGT are circled in dotted lines. Adapted from “Custom Plasmid Maps 2”, by BioRender.com (2022). Retrieved from https://app.biorender.com/biorender-templates

### Deletion of *craA*

For the first trial of this genome editing method, we attempted the construction of an in-frame deletion mutant in *craA* (ABUW_0337). A visual outline of the strategy can be followed in Figure 2. Once the pEMGT derivative bearing the flanking homologous regions of *craA* was constructed (pEMGT-*craA*), it was conjugated into the AB5075 parental strain and transconjugants bearing the plasmid inserted by recombination were selected in the presence of tellurite. Five candidates were confirmed to carry the plasmid integrated into the chromosome by PCR (Supplementary Figure S1), and transconjugants appeared with frequency of 10^−8^.

**Figure 2.**
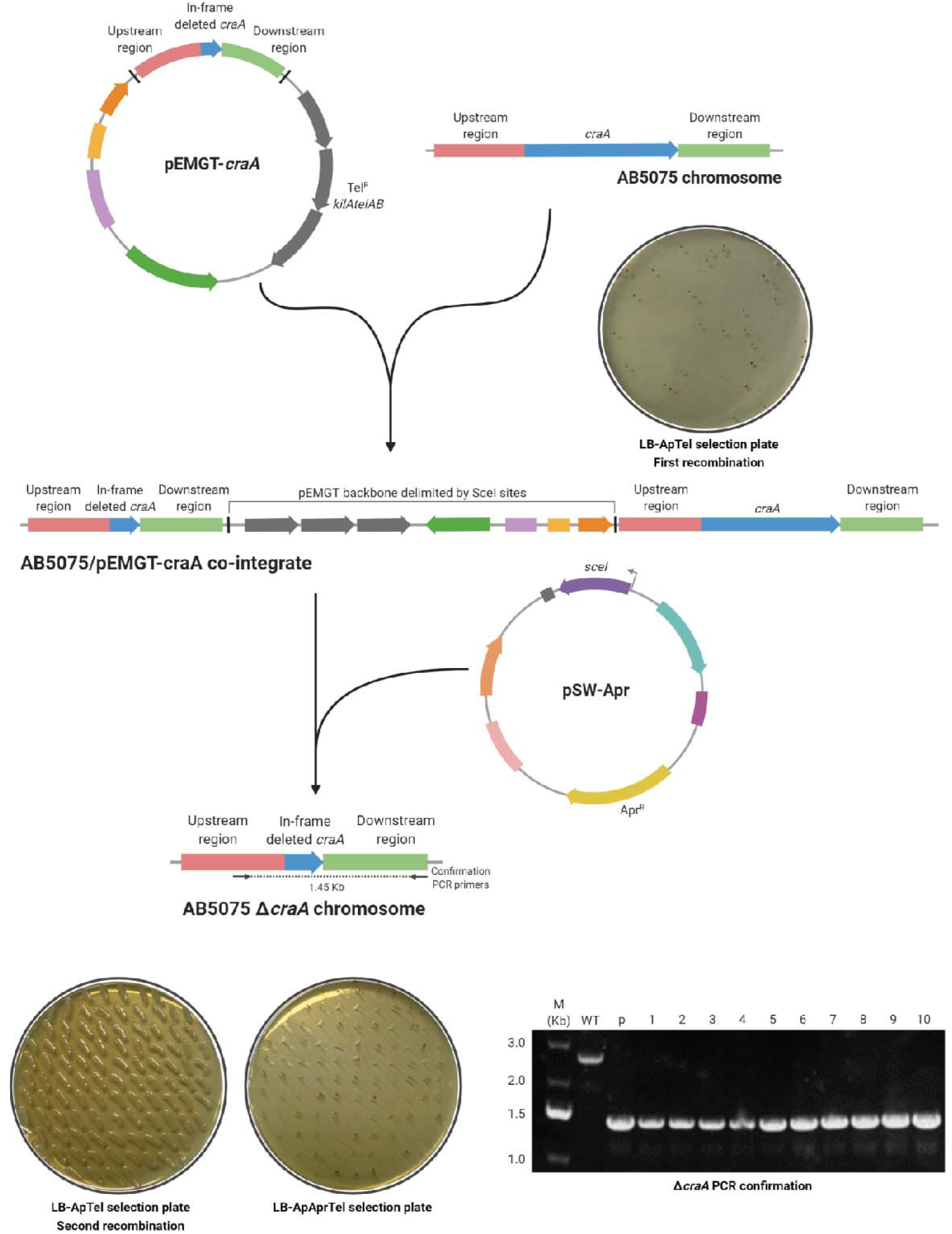
Schematic outline of the genome editing strategy adapted for *A. baumannii* AB5075 applied to the deletion of *craA*. Plasmid features are represented as in Fig. 1. When indicated, LB agar plates were supplemented with ampicillin 100 mg/L (Ap), apramycin 200 mg/L (Apr) and/or tellurite 30 mg/L (Tel). For confirmation of *craA* deletion, colony PCR was performed using primers craA fw seq and craA down rv. As controls, wild type AB5075 (WT) and pEMGT-*craA* (p) were used. M: DNA molecular weight marker, with band sizes indicated in kilobases (Kb). Created with BioRender.com.

Among the candidates, three colonies were selected for performing the second recombination event. To check the effectiveness of both pSW-Apr and pSW-Tc for forcing the second recombination event, both of them were transferred by mating in biological triplicates to the AB5075-pEMGT-*craA* parental strain and transconjugants were selected in the presence of either antibiotic. We attempted the on-plate *sceI* induction by adding the inducer 3-methylbenzoate (3MB) to the selective plates. However, the presence of this compound affected *A. baumannii* growth (Supplementary Figure S2). Nevertheless, this strategy has been applied before without addition of the inducer,^30-32^ which also resulted successful for *A. baumannii*. In the case of pSW-Apr recipients, clear individual colonies grew with a frequency around 10^−4^. However, although pSW-Tc recipients grew with a similar frequency, colonies appeared with a viscous, squashed phenotype (which we had previously observed when selecting tetracycline resistance) that madeselection difficult (Supplementary Figure S3).

To assess the second recombination, we screened for the loss of tellurite resistance. This screening resulted in 98.0 ± 1.7 % of clones that achieved a second recombination triggered by presence of pSW-Apr (Figure 2) and 72.3 ± 3.2 % of clones by pSW-Tc.

To select a double recombinant carrying the in-frame deletion of *craA* instead of a reversion to wild type genotype, 10 random candidates among all the pSW-Apr transconjugants were streaked to obtain individual colonies and analysed by PCR. In this case, the screening resulted in 100% deletion frequency according to the size of the PCR product.

As a final step in the protocol, the resulting mutant strain had to be cured from pSW-Apr. For this, one mutant clone was inoculated in LB broth in the absence of apramycin and two passages were given after reaching saturation. After this, individual colonies were isolated and screened for apramycin sensitive clones. Chromosomal deletion was checked by sequencing (Supplementary Figure S4, Supplementary File S1). To facilitate use of this strategy, a detailed step-by-step laboratory protocol in 7-9 days is shown as Supplementary Text S1.

### Deletion of *cmlA5* and removal of p1AB5075

In order to know if this mutagenesis system would be suitable for native plasmid editing within *A. baumannii*, we challenged it by attempting the deletion of *cmlA5* (*ABUW_4059*). This gene encodes a putative chloramphenicol efflux pump and is located within the so-called resistance island 2 (RI2), borne in the p1AB5075 plasmid.^22,33,34^

For the deletion, we performed a similar strategy as for the mutation of *craA*. Once the respective flanking homologous regions were cloned into pEMGT (pEMGT-*cmlA5*), the plasmid was transferred to AB5075 and its integration was selected for. For the second recombination, we leaned toward using pSW-Apr, given its better performance compared to pSW-Tc. After screening for a second recombination event, we checked 20 candidates by PCR. In this particular case, we found that, whereas 35% of the clones had suffered a second recombination (they gave a PCR of either wild type or mutant size), the remaining 65% did not yield any amplification product. This would indicate that either a rearrangement in the plasmid had occurred, removing the region that served as PCR template, or that the whole plasmid had been removed. A possible explanation would be a scission of RI2 by homologous recombination between the two miniature inverted-repeat transposable element (MITE)-like sequences that flank this island, explaining the loss of the *cmlA5* region while keeping the rest of p1AB5075.^22,35^ To address this, the same 20 candidates were analysed by PCR with two primer pairs that had been used previously to check the presence of p1AB5075.^36^ This resulted in 20% of them not yielding a PCR product with any primer pair, indicating the loss of the plasmid. One example of each template-primer pair combination is shown in Figure 3.

**Figure 3.**
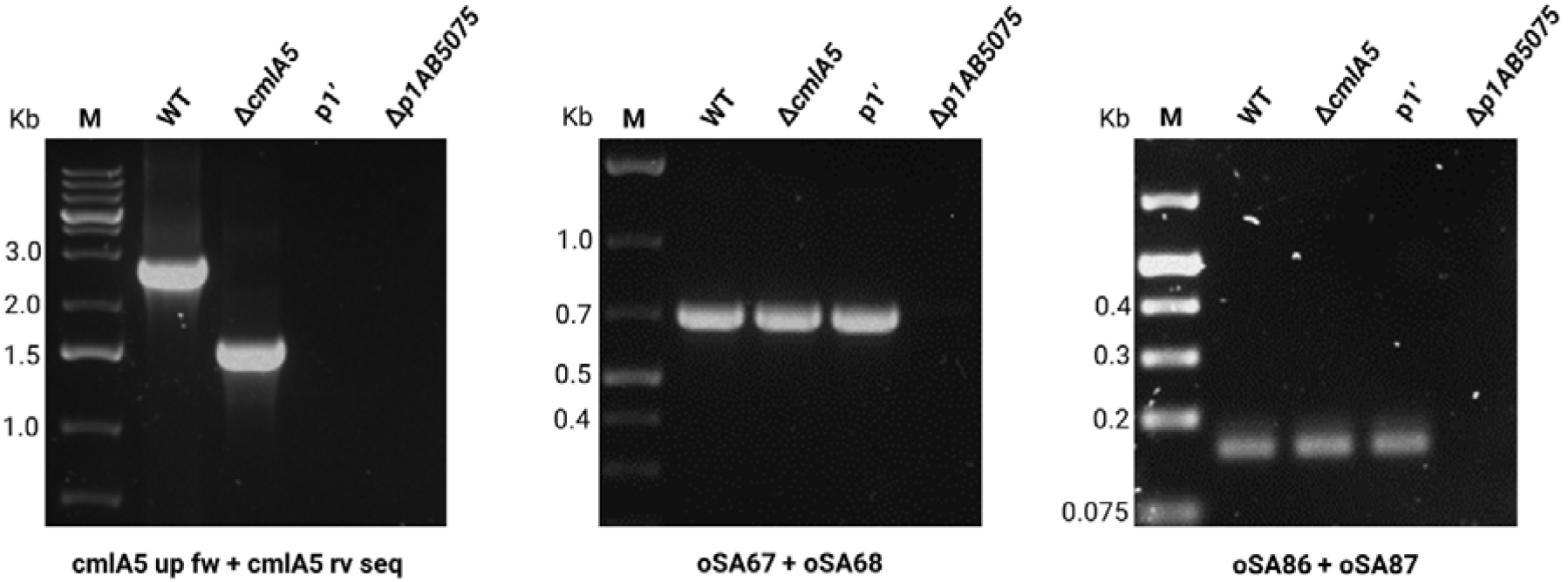
PCR analysis to confirm the Δ*cmlA5* deletion and assess the presence of p1AB5075. Genomic DNA extracted from the respective strain was used as template. For confirming the deletion, primer pair cmlA5 up fw/cmlA5 rv seq was used, giving bands of 2.79 Kb for the wild type (WT) and 1.55 Kb for the Δ*cmlA5* deletion mutant. The presence or absence of p1AB5075 was assessed with primer pairs oSA67/oSA68 and oSA86/oSA87, which would give PCR products of 0.7 Kb and 0.16 Kb, respectively (Anderson 2020). In the case of the p1AB5075-cured strain, no amplification was observed for any of the primer pairs. A plasmid rearrangement in p1AB5075 (p1’) is suggested by the absence of PCR product using the primers to detect *cmlA5* and compared to the amplification with primers to confirm presence of p1AB5075. M: DNA molecular weight marker, with band sizes indicated in Kb.

Curing native plasmids, usually of unknown function, often involves tedious counterselection screenings.^37-39^ Else, spontaneous plasmid-cured strains can be found serendipitously.^36,40^ Given the high frequency of *A. baumannii* strains bearing multiple native plasmids and the difficulties entailed by mutating and manipulating them, this methodology shows a remarkable potential for their study.

### Phenotypic characterisation of the *craA* and *cmlA5* mutants

To assess the efficiency of this mutagenesis method, we chose to delete genes encoding for antibiotic resistance, whose phenotype can be measured easily. Whereas there are reports about the function of CraA,^28,29^ the RI2-encoded CmlA5 has only been annotated as a chloramphenicol efflux pump based on homology.^22,33,34^ The phenotypic characterisation and comparison of both mutants would help us elucidate the contribution of CmlA5 to antibiotic resistance in AB5075.

For their characterisation, we assessed chloramphenicol resistance by disc diffusion assays (DDA) and minimum inhibitory concentration (MIC) measurements comparing both deletion mutants, as well as the p1AB5075-cured strain, to the wild type (Figure 4, Table 1, Supplementary Figure S5). Whereas the DDAs showed that the Δ*craA* mutant was the only one with an increased sensitivity to chloramphenicol (3.125 mg/L compared to 200 mg/L for AB5075), the MIC assays revealed a mild increase in sensitivity for the Δ*cmlA5* mutant, that kept increasing for the p1AB5075-cured strain (100 mg/L and 50 mg/L, respectively). This indicates a minor role of *cmlA5* in chloramphenicol resistance in AB5075 and suggests there are additional chloramphenicol resistance determinants encoded in p1AB5075.

**Table 1.** Minimum inhibitory concentration (MIC) for the Δ*craA* and Δ*cmlA5* mutants and the p1AB5075-cured strain (Δ*p1AB5075*) compared to the wild type AB5075. MICs were assessed in CAMH broth. Antibiotics were diluted by 2-fold, ranging from 400 mg/L to 0.781 mg/L, except for tobramycin when used against the Δ*craA* mutant, which ranged from 100 mg/L to 0.195 mg/L. The MIC was assessed as the first concentration that showed no visual growth. Three biological replicates were conducted.

| | AB5075 | ΔcraA | ΔcmlA5 | Δp1AB5075 |
| --- | --- | --- | --- | --- |
| **Chloramphenicol** | 200 mg/L | 3.125 mg/L | 100 mg/L | 50 mg/L |
| **Amikacin** | 200 mg/L | 1.5625 mg/L | 200 mg/L | 3.125 mg/L |
| **Tobramycin** | 25 mg/L | 1.5625 mg/L | 25 mg/L | < 0.781 mg/L |

**Figure 4.**
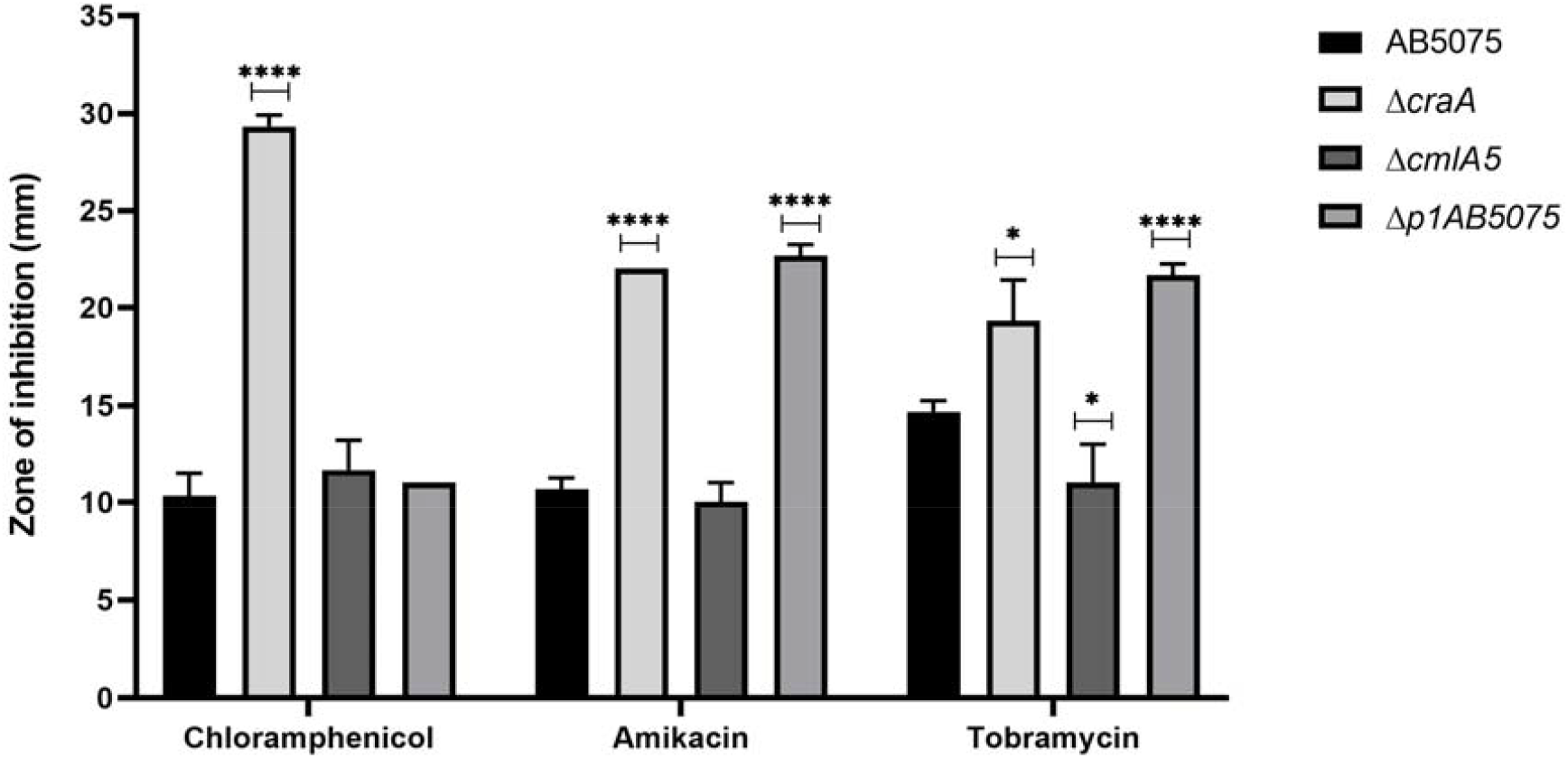
Antibiotic disc diffusion assay for the Δ*craA* and Δ*cmlA5* mutants and p1AB5075-cured strain compared to the wild type AB5075. Assays were performed in CAMH agar assessing sensitivity to imipenem (10 μg) as control (no increase in sensitivity was expected for imipenem in these strains compared to AB5075), chloramphenicol (50 μg), amikacin (30 μg) and tobramycin (10 μg). The average zone of inhibition in millimetres (mm) measured from 3 biological replicates ± S.D. is shown. Statistical significance was assessed from P-values obtained from a t-test (* = P ≤0.05, ** = P ≤0.01, **** = P ≤0.0001).

Since *cmlA5* is located within RI2, likely forming an operon with genes related to aminoglycoside resistance (*aadA1, strA, strB*), we wondered whether it could play a role in resistance to this class of antibiotics.^22,36^ To address this, we performed DDAs and MIC assays for the aminoglycosides amikacin and tobramycin for the two mutants and the p1AB5075-cured strain compared to AB5075 (Figure 4, Table 1, Supplementary Figure S5). Interestingly, whereas both DDAs and MICs showed that *cmlA5* is unrelated to aminoglycoside resistance (same MICs for Δ*cmlA5* and AB5075), they showed that CraA confers resistance to amikacin and tobramycin, with MICs dropping more than 130-fold. As expected, the loss of p1AB5075 led to a substantial reduction in aminoglycoside resistance, particularly to tobramycin. It was also worth mentioning a slight increase in resistance to tobramycin for the Δ*cmlA5* mutant. Since *cmlA5* is likely the first of an operon formed with other three genes related to antibiotic resistance, it would be possible that its deletion minorly altered the expression of the following genes, thus leading to this mild increase.

Although CraA was previously thought to confer resistance specifically to chloramphenicol,^28^ it was recently shown to have a broader substrate range.^29^ Consequently, it was postulated to be closer in function to the multidrug efflux pump MdfA, although differing in the substrate recognition mechanism.^29^ Here, we expand the CraA substrate spectrum even further by demonstrating its role in aminoglycoside resistance. Altogether, this indicates that CraA might play a major role in *A. baumannii* multidrug resistance.

## Acknowledgements

The authors would like to thank Dr Esteban Martínez-García and Prof Víctor de Lorenzo (Centro Nacional de Biotecnología, Madrid, Spain) for providing pEMG, pSW-I and pSEVA524 as a kind gift. RRMC is supported by the British Society for Antimicrobial Chemotherapy BSAC-2018-0095. RRMC and RD are supported by a BBSRC New Investigator Award BB/V007823/1. RRMC is supported by the Academy of Medical Sciences/the Wellcome Trust/ the Government Department of Business, Energy and Industrial Strategy/the British Heart Foundation/Diabetes UK Springboard Award [SBF006\1040].

## Author contribution

RD and RRMC designed the strategy and the experimental work. RD and KG performed the experiments and analysed the results. RD, KG and RRMC wrote and reviewed the manuscript.

## Data Availability

All plasmids available through request to the corresponding author.

## Transparency declarations

The authors declare no competing interests.

