## Supplementary Information for "A high-efficiency scar-free genome editing toolkit for *Acinetobacter baumannii*"

Rubén de Dios[1], Kavita Gadar[1] and Ronan R McCarthy[1]

[1] Division of Biosciences, Department of Life Sciences, College of Health and Life Sciences, Brunel University London, Uxbridge, UB8 3PH, UK.

Running title: Scar-free mutagenesis in *Acinetobacter baumannii*

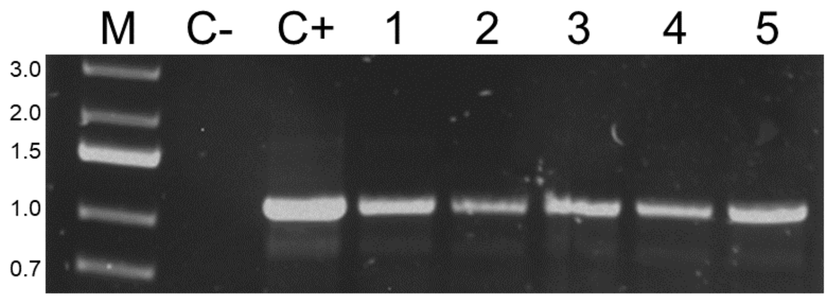


**Supplementary Figure S1. PCR checking for the integration of pEMGT-*craA* into the AB5075 chromosome.** The primer pair used was M13 rev (anneals in pEMGT flanking the polylinker) and craA up rv (anneals within the craA deletion construct). As negative control (C-) we used 20 ng of AB5075 genomic DNA; as positive control (C+) we used pEMGT-*craA*. Co-integration candidates are numbered 1-5. M: DNA molecular weight marker, with band sizes on the left in Kb.

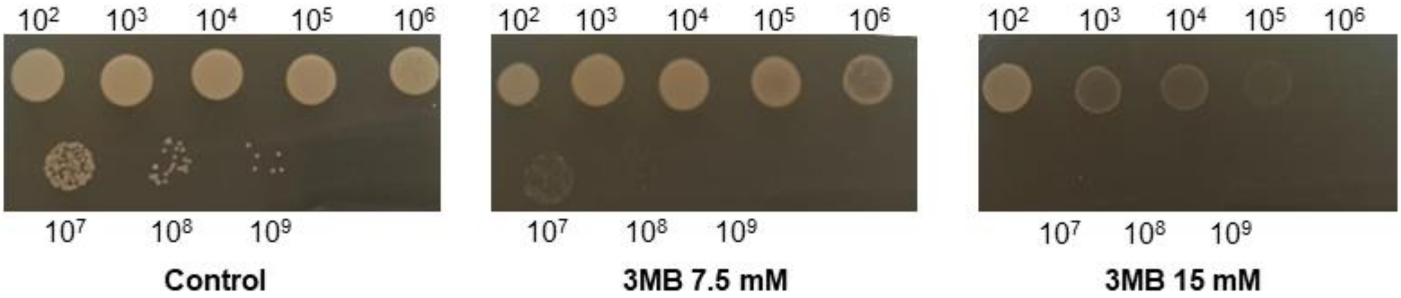


**Supplementary Figure S2. Inhibitory effect of 3-methylbenzoate (3MB) on *A. baumannii* AB5075.** A overnight culture of AB5075 was serially diluted (dilution factor indicated at the top and the bottom) and 10 µl of each dilution were spotted on plain LB agar (Control) or agar supplemented with 3MB 7.5 mM or 15 mM (concentration used in Martínez-García and de Lorenzo, 2011). Plates were incubated at 37 C overnight. Both 3MB concentrations inhibit the growth of AB5075.

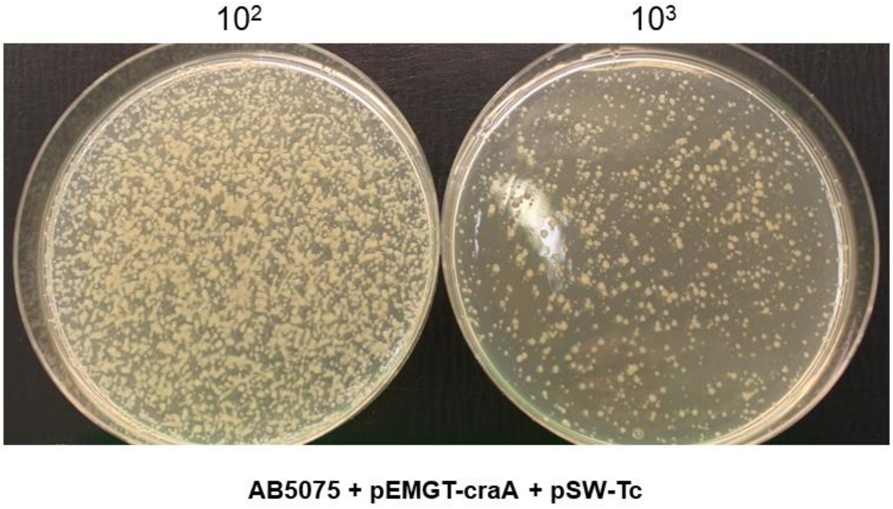


**Supplementary Figure S3. Colony phenotype of *A. baumannii* AB5075 in the presence of tetracycline (Tc).** After conjugation of pSW-Tc into the AB5075/pEMGT-*craA* co-integrate strain, the biomass was resuspended in 1 ml LB and serial dilutions were plated in LB agar supplemented with Tc (5 mg/L). Plates spread with dilution factor $10^2$ and $10^3$ are shown. Colonies grew exhibiting a viscous, squashed phenotype that hindered the candidate selection.

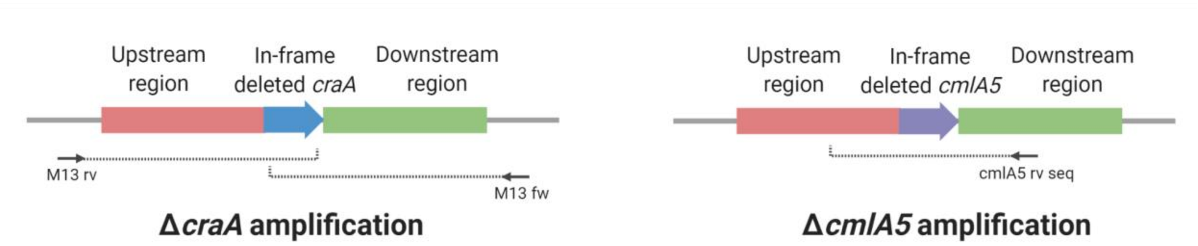


**Supplementary Figure S4. Schematic of the sequencing performed for confirmation of the Δ*craA* and Δ*cmlA5* mutants.** After PCR confirmation of clones bearing the Δ*craA* and Δ*cmlA5* deletions, the genomic DNA of each was PCR amplified using primer pairs craA up fw/craA down rv and cmlA5 up fw/cmlA5 down rv, respectively. The PCR products were sub-cloned into pEMG digested with SmaI and sent for sequencing using primers M13 fw and M13 rv for the Δ*craA* amplification and cmlA5 rv seq for the Δ*cmlA5* amplification. The reach of each sequencing reaction is indicated with dotted lines, and the primers are indicated with arrows. Chromatogrammes resulting from the sequencing reactions are provided in Supplementary File S1.

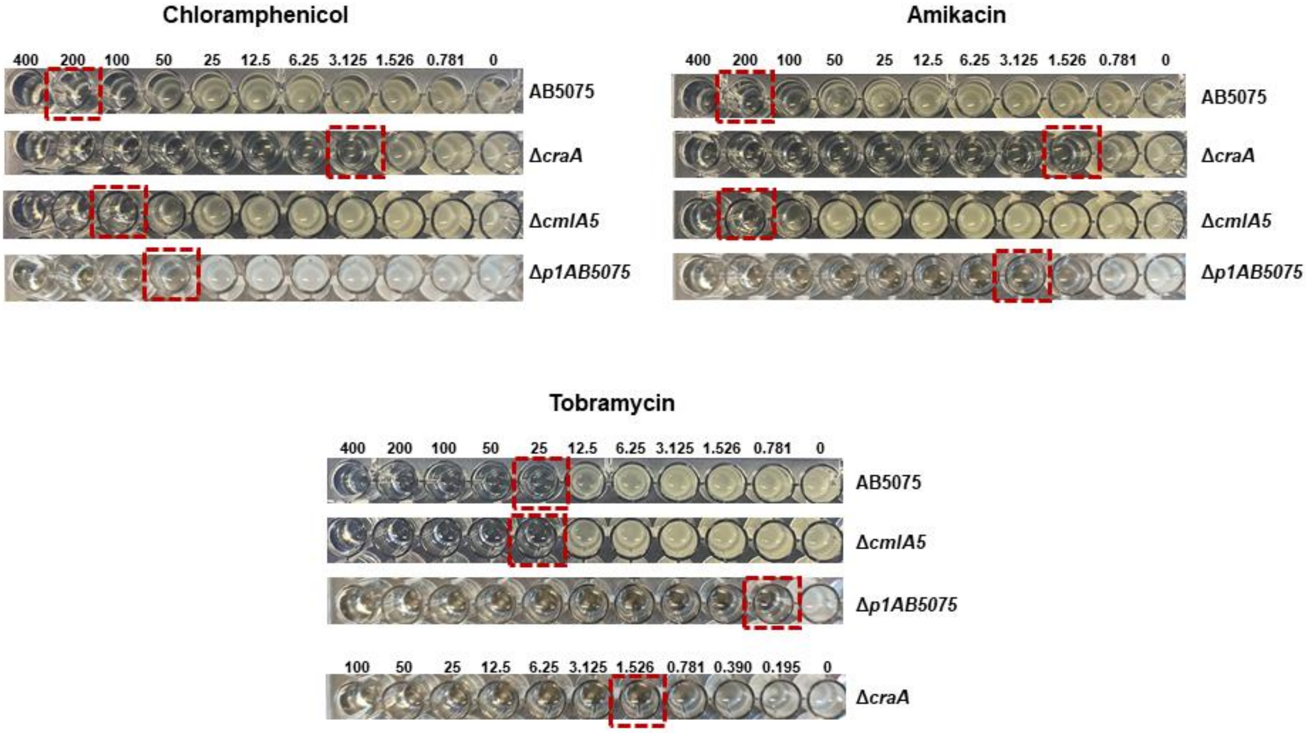


**Supplementary Figure S5. Visual examination of minimum inhibitory concentration (MIC) for chloramphenicol, amikacin and tobramycin.** MICs were assessed for AB5075, the Δ*craA* and Δ*cmlA5* mutants and the p1AB5075-cured strain (Δ*p1AB5075*) as the first antibiotic concentration that produced no visual growth (highlighted in red). Antibiotic concentrations are indicated above each set of rows in mg/L. Antibiotic 2-fold dilutions ranging from 400 mg/L to 0.781 mg/L were used in all cases, except for tobramycin with the Δ*craA* mutant, for which the range was 100 mg/L to 0.195 mg/L. The tobramycin MIC for the p1AB5075-cured strain was assessed as < 0.781 mg/L. Representative images out of 3 biological replicates are shown.

**Supplementary File S1. Chromatogrammes obtained from sequencing the genomic regions around the Δ*craA* and Δ*cmlA5* deletions.** Genomic DNA from each mutant was amplified with primer pairs craA up fw/craA down rv and cmlA5 up fw/cmlA5 down rv, respectively. The PCR products were sub-cloned into pEMG digested with SmaI and sent out for sequencing using primers M13 fw and M13 rv for the Δ*craA* amplification and cmlA5 rv seq for the Δ*cmlA5* amplification.

**Supplementary Table S1. Bacterial strains, plasmids and PCR primers used in this work.** Attached as three different tabs in an Excel file.

**Supplementary Text S1**

**Standard protocol for genome editing in *A. baumannii* AB5075**

**All plasmids for this protocol are available for free by request to the McCarthy Lab.**

Using pEMGT and pSW-Apr

Time required: 7 - 9 days

Material (glassware and plasticware not included)

- LB agar and broth
- Antibiotic stocks: tetracycline 5 mg/mL (Tc), ampicillin 100 mg/mL (Ap), kanamycin 25 mg/mL (Km), apramycin 60 mg/mL (Apr60, for *E. coli*) and 200 mg/mL (Apr 200, for *A. baumannii*)
- Potassium tellurite 30 mg/mL (Tel)
- pEMGT-derivative in an *E. coli* DH5α λ*pir* host
- pSW-Apr in an *E. coli* DH5α host
- pRK2013 helper plasmid in an *E. coli* DH5α host (any other helper plasmid suitable for triparental mating should work)
- DreamTaq Green PCR Master Mix (ThermoFisher)
- Appropriate primer pair for clone confirmation
- Agarose and gel casting, running and visualization equipment

Day 0

1. Inoculate AB5075, pRK2013 and the pEMGT-derivative in 5 ml LB broth supplemented with Ap for AB5075 and Km for the other two cultures. Incubate overnight at 37 ºC, 180 rpm.

Day 1

1. Mix 500 µl of AB5075, pRK2013 and the pEMGT-derivative in a 1.5 ml tube. Centrifuge maximum speed for 30 s.
2. Discard the supernatant and wash 2 times with fresh LB broth.
3. After the last wash, resuspend the pellet in 40 µl of LB broth and spot the mixture on an LB agar plate.
4. After air-drying, incubate at 37 ºC for 4 h.
5. Resuspend the patch in 1 ml LB broth.

6. Plate dilutions with dilution factors $10^0$-$10^1$ on LB agar supplemented with Ap and Tel. Incubate overnight at 37 ºC.

Day 2

1. Transconjugants must have appeared with a frequency of $10^{-8}$. Mark them with numbers in the selection plate and take samples to check by PCR. Note: tellurite is a strong selection, if plates are not over-incubated, all transconjugants must bear the plasmid insertion. Thus, the PCR confirmation may be avoided.
2. Inoculate the selected candidate in LB broth (no selection needed, the insertion is stable enough). Also, inoculate pRK2013 and pSW-Apr in LB broth supplemented with Km and Apr60, respectively. Incubate overnight at 37 ºC, 180 rpm.

Day 3

1. Mix 500 µl of AB5075, pRK2013 and pSW-Apr in a 1.5 ml tube. Centrifuge maximum speed for 30 s.
2. Discard the supernatant and wash 2 times with fresh LB broth.
3. After the last wash, resuspend the pellet in 40 µl of LB broth and spot the mixture on an LB agar plate.
4. After air-drying, incubate at 37 ºC for 4 h.
5. Resuspend the patch in 1 ml LB broth.
6. Plate dilutions with dilution factors $10^2$-$10^4$ on LB agar supplemented with Ap and Apr200. Incubate overnight at 37 ºC.

   Note: To accelerate the protocol, Day 2 and Day 3 can be performed in one day avoiding the PCR confirmation. pRK2013 and pSW-Apr must be inoculated on Day 1 and the selected AB5075/pEMGT-derivative clone must be inoculated in LB broth early in the morning. After approx. 7 h, there should be enough biomass for performing the mating as in Day 3.

Day 4

We have observed that triggering the double recombination on the selection plate can lead to mixed population colonies, with some of the cells having undergone the second recombination and some others not. If candidates are streaked directly from here as in Figure 2 (main text) and are checked by PCR, there is the possibility of obtaining two amplifications with the sizes of the wild type and the mutant. To avoid this, we strongly recommend segregating the colonies to obtain isolated clones from them either before or after a marker exchange screening.

1. Streaking candidates in LB agar supplemented with Apr200 to obtain isolated colonies. Incubate overnight at 37 ºC.

Day 5

1. Double-streak selected clones on LB agar supplemented with Tel and Apr200 and Apr200 only to check if they lost the Tel resistance. Incubate overnight at 37 ºC.

Day 6

1. Select the candidates that grew on LB with Apr200 but not on LB with Tel and Apr200. Select 5-10 for further screening.
2. Screen the selected candidates by PCR (using the appropriate primer pair) to confirm that the genome modification has been performed successfully.
3. Select one clone that both lost the Tel resistance and gave the appropriate PCR product size. Further confirmation can be done by sequencing the region on the modification.
4. Inoculate the selected candidate in plain LB broth to cure pSW-Apr. Incubate overnight at 37 ºC, 180 rpm.

   Note: To accelerate the protocol, Day 5 and Day 6 can be performed in one day. If Day 5 streaking is performed early in the morning, after 7 hour incubation there should be enough biomass in the streaks to confirm the marker exchange and use it as a PCR template. After knowing the result of the PCR, the appropriate candidate can be inoculated to start the curation of pSW-Apr.

Day 7

1. Dilute the overnight culture in fresh LB broth 1:1000. Incubate at 37 ºC, 180 rpm for 10 h.
2. Streak 10 µl on an LB agar plate to obtain isolated colonies. Incubate overnight at 37 ºC.

   Note: To ensure the loss of pSW-Apr, an additional passage can be performed. However, at least for AB5075, we have observed that 3 passages increase the rate of phase variation, requiring further selection of the colony morphotype in the next step.

Day 8

1. Double-streak the isolated colonies cured from pSW-Apr on plain LB agar and on LB agar supplemented with Apr200. Incubate overnight at 37 ºC.

Day 9

1. Select a clone that grew on plain LB but not on LB supplemented with Apr200.
2. Unless further confirmation is required, the genome-edited *A. baumannii* strain is ready to stock and use for experiments.
