## Supplementary Table S1 for "A high-efficiency scar-free genome editing toolkit for *Acinetobacter baumannii*"

| **Strain** | **Relevant features** | **Reference** |
|---|---|---|
| *Escherichia coli* | | |
| DH5α | F- ϕ80lacZΔM15, Δ(lacZYA-argF)U169, recA1, endA1, hsdR17(rK-m K-), supE44, thi-1, gyrA, relA1 | Hanahan, 1983 |
| DH5α*λpir* | F- ϕ80lacZΔM15, Δ(lacZYA-argF)U169, recA1, endA1, hsdR17(rK-m K-), supE44, thi-1, gyrA, relA1, λpir lysogen | Martínez-García & de Lorenzo, 2011 |
| *Acinetobacter baumannii* | | |
| AB5075 | Wild type strain | Jacobs *et al*., 2014 |
| Δ*cmlA5* | AB5075 derivative, in-frame deletion in *cmlA5* (ABUW_4059) | This work |
| Δ*craA* | AB5075 derivative, in-frame deletion in *craA* (ABUW_0337) | This work |
| p1' | AB5075 derivative, indeterminate rearrangement in p1AB5075 loosing *cmlA5* region (likely the whole resistance island 2) | This work |
| Δ*p1AB5075* | AB5075 derivative, cured from p1AB5075 plasmid | This work |
| **Plasmids** | **Relevant features** | **Reference** |
| pEMG | KmR, *ori*R6K, *lacZa* with two flanking I-SceI sites | Martínez-García & de Lorenzo, 2011 |
| pEMGT-*cmlA5* | pEMGT with upstream and downstream homologous regions for deletion of *cmlA5* | This work |
| pEMGT-*craA* | pEMGT with upstream and downstream homologous regions for deletion of *craA* | This work |
| pEMG-Tel (pEMGT) | KmR, TelR, *ori*R6K, *lacZa* with two flanking I-SceI sites | This work |
| pFLAG-attP | AprR, CmR, *attP* integration site (Addgene, #110095) | Unpublished |
| pMo130-TelR | KmR, TelR, *sacB*, *xylE* | Amin *et al*., 2013 |
| pRK2013 | KmR, ColE1, tra+. Conjugative helper plasmid. | Figurski and Helinski, 1979 |
| pSEVA524 | TcR, oriRK2, *lacIq-Ptrc* | Silva-Rocha *et al*., 2013 |
| pSW-Apr | AprR, *ori*RK2, *xylS*, Pm→*sceI* (transcriptional fusion of I-*sceI* to *Pm*) | This work |
| pSW-I | ApR, *ori*RK2, *xylS*, Pm→*sceI* (transcriptional fusion of I-*sceI* to *Pm*) | Martínez-García & de Lorenzo, 2011 |
| pSW-Tc | TcR, *ori*RK2, *xylS*, Pm→*sceI* (transcriptional fusion of I-*sceI* to *Pm*) | This work |
| **Primer** | **Sequence** | **Reference** |
| Apr fw | AGAGGAGACCTAGTTGGCAC | This work |
| Apr rv | GAGCTGAAGAAAGACAATCC | This work |
| cmlA5 down fw | GATGGTTTCGTGCGCTCAAAACGTTGAGAGAATGTGGCAAG | This work |
| cmlA5 down rv | GGAAGCAGGTGCGATTTTCG | This work |
| cmlA5 rv seq | ATCAATGTCGATCATGGCTG | This work |
| cmlA5 up fw | AGTTGACATAAGCCTGTTCGG | This work |
| cmlA5 up rv | TTTTGAGCGCACGAAACCATC | This work |
| crA up fw | CATTTAATTCACCATTTGAG | This work |
| craA down fw | GAAGTGTACAAACTCATGAAAAGAGTTGCACAAGACTTAC | This work |
| craA down rv | CTATTTAGACATGCAGTACG | This work |

Supplementary Table S1

| | | |
|---|---|---|
| craA fw seq | CGAGGAAGCAGCAGACTTTC | This work |
| craA up rv | TTCATGAGTTTGTACACTTC | This work |
| oSA67 | GACGCCCGTCTAACAATTCG | Anderson *et al.*, 2020 |
| oSA68 | CCCCTCGATGGAAGGGTTA | Anderson *et al.*, 2020 |
| oSA86 | TAAGCGTCAGGCAGACAAG | Anderson *et al.*, 2020 |
| oSA87 | TTTTCCACTCTGCTGAAGG | Anderson *et al.*, 2020 |
| tetA fw | TAAATTTGACAGCTTATCATCG | This work |
| tetA rv | AAAGGACAATTGTCTCAGGTCG | This work |
